# Water yield modelling, sensitivity analysis and validation: a study for Portugal

**DOI:** 10.1101/2021.03.20.433030

**Authors:** Bruna Almeida, Pedro Cabral

**Affiliations:** NOVA Information Management School (NOVA IMS), Universidade Nova de Lisboa, 1070-312 Lisboa, Portugal

## Abstract

The spatially explicit assessment of freshwater is key to introduce the ecosystem service (ES) concept into decision-making processes. Many tools are being developed to model water balance and to analyse the effects of meteorological conditions on water ES behaviours at multiple spatial scales. Nevertheless, there is a lack of studies that analyse the sensitivity of these models and estimate the model’s accuracy. The current research uses the InVEST Water Yield Model (WYM) to assess freshwater ES between 1990 and 2018 at River Basin District (RBD) level in mainland Portugal. The methodology included sensitivity analysis simulations to test different parameters of the model over two different sources of meteorological data. We validated the model with the European Environment Agency (EEA) database on the quantity of Europe’s freshwater resources. To evaluate the models’ sensitivity, Pearson’s correlation coefficients and statistical methods were calculated for each simulation. The model which obtained the best performance in the sensitivity tests was retained for further analysis and calibration and its accuracy was assessed by comparing the mean estimated water yield (WY) with mean observed values for 2018. Results at the national level show a correlation coefficient of 0.803 with statistical significance for 0.01 one-tail. The WY in the RBDs of the North of the country was underestimated by 56.5 mm/ha/year and for the RBDs in the South of Portugal, the WY was overestimated by 58.1 mm/ha/year. This difference was explained through the spatial-temporal assessment of the main climatic variables used as input. This study contributes to a methodology to assess the level of confidence in WYM outputs and can be used to support the trustworthiness of water availability studies at sub-watershed, watershed, or RBD, using open access data and software.

## 1. Introduction

Ecosystem services (ES) represent the benefits human populations derive, directly or indirectly, from ecosystem functions [1]. Most of nature’s contributions to people are not fully replaceable, and some are irreplaceable [2]. The long-term security of many ecosystem functions and services, especially in changing environments, is likely to depend upon local biodiversity, which has been facing growing pressures from human activities [3]. Some of those activities result in modifications in the natural landscapes around rivers and wetlands to the point that their biodiversity is put at risk [4]. Freshwater ecosystems are crucially important providers of ecosystem services as they provide clean water for multiple uses including water for drinking, irrigation, or recreation [5]. Since 1970, land-use change has had the largest relative negative impact on this service, followed by direct exploitation, in particular, overexploitation [2].

Climate change poses another set of pressures on water-related ES [6] with extreme weather events perceived as the top risk affecting immediately the agricultural sector [7]. Over one-third of the world’s land surface and nearly three-quarters of available freshwater resources are devoted to crop or livestock production [2]. The water demand is strongly influenced by population and agricultural production, and water availability is linked to the frequency and amount of precipitation [8]. The negative impacts of climate change on water availability call for urgent action to allocate and to use water more wisely [6]. More than half the world’s population, some four billion people, are exposed to drought for at least one month per year [7].

For the seventh consecutive year, the Global Risks Report 2018 listed water crises among the top five global risks both in terms of impact and in terms of likelihood [9]. To deal with that, adaptive management practices must be implemented to increase the adaptation capacity of river basins to changing conditions, such as climate and global change [5]. River Basin District (RBD) represents the natural boundaries of a complex ecological system [10] that according to the report carried out by the Water Resources Group in 2009, by 2030 over a third of the world population will be living in river basins that will have to cope with significant water stress [11].

Projections indicate that the most rapid increase in water demand will be in urban contexts [7]. As climate change and the lack of adequate adaptation strategies are the biggest threat to Europe’s water environment [12], an approach to face this issue is to identify sustainable pathways that would anticipate and manage the inevitable increases in water scarcity and water-related risks [6]. River basins resilience measure the degree to which a catchment can adjust to normal functioning following perturbations from events such as drought or extreme precipitation [13] and also under some activities such as energy production and crop irrigations [12].

A step towards the harmonisation of water, ecosystems and society can be achieved through the assessment, mapping and valuation of ecosystem services [5]. Nowadays, we are seeing a more explicit quantification of the value of nature in defining human-water interactions [4]. Several spatially-based decision support tools have emerged for ESs assessment [14], and one of them is the InVEST (Integrated Valuation of Ecosystem Services and Tradeoffs) developed by the Natural Capital Project - Stanford University. The InVEST Reservoir Hydropower Production Water Yield Model (WYM) estimates the relative contributions of water from different parts of a landscape, offering insights into how changes in land-use patterns affect water balance, freshwater provision accounts and hydropower production [15].

The mapping and evaluation of ecosystems and their services is a key action to communicate the characteristics, trends, and rate of change in ecosystem services and biodiversity [16]. It provides useful information for decision-makers [17] in planning and acting to strengthen the resilience of economies and societies to protect them from water-related disasters [6]. The existing challenges to introduce the ESs concept into decision-making processes are related to the requirement for accurate mapping and evaluation, and a deep understanding of models’ sensitivities [15].

Sensitivity analysis is an effective tool for identifying important uncertainty sources and improving model calibration and predictions [18]. It is an integral part of model development and involves an analytical examination of input parameters to aid in model validation and provide guidance for future research [19]. This technique consists of changing values of inputs and internal parameters of a model to determine the effect on the behavior and output of the model [20], increasing our understanding of which parameters must be determined with the greatest accuracy, and thus helping to prioritize data collection requirements [18].

All parameters in the InVEST WYM display some degree of sensitivity [21]. Previous studies (e.g. [22]) show that the model can be very sensitive to changes in drivers such as land-use and meteorological data. Extremes variations in those drivers can alter hydrologic cycles, affecting patterns of evapotranspiration, infiltration, and water retention [23]. The goal of the InVEST model is not to reproduce observations with a high degree of accuracy and precision, but to provide reliable geospatial information to support decisions [24]. However, an important and often overlooked question is how accurate are the outputs of the WYM with empirical observations [15].

This paper aims to answer this question by studying the WYM uncertainties and estimating its accuracy, through sensitivity analysis and validation. The model’s sensitivity is evaluated by testing: two different sources of climatic data; different values for the seasonality coefficient (Z); different grids of plant available water content and biophysical table, and model’s outputs were validated with the European Environment Agency (EEA) database on the quantity of Europe’s water resources of 2018. Model’s accuracy was calculated comparing mean estimated water yield with mean observed values from the validation data, and to evaluate the model’s sensitivity, Pearson’s correlation coefficients and statistical methods were calculated for each simulation. Analysing the sensitivity of the WYM and estimating its accuracy aim to build confidence in the model’s outputs, and to support studies related to the spatial estimation of water availability and water ESs’ behaviours at sub-watershed, watershed, or RBD.

## 2. Materials and methods

### 2.1. Study area

Portugal is in the Iberian Peninsula (south-western Europe) comprising a continental part and two autonomous regions, Azores, and Madeira archipelagos. The territory total area is 92 072 km^2^ and the population is about 9.8 million [25].

In the climatic perspective, it lays in the transitional region between the subtropical anticyclone and the sub-polar depression zones [26]. The most conditioning climate factors are latitude, orography and the effect of the Atlantic Ocean, characterizing high spatial and temporal variability [27] that affect the water cycle primarily through changes in the precipitation and temperature [28]. In mainland Portugal, the large temporal variability of the precipitation rates leads to precipitation extreme events and intense dry months, resulting in impacts in water resources, fire risk and ecosystem degradation [27].

As other southern European regions, Portugal is quite vulnerable to climate variability, namely to droughts and desertification, especially in the southern sector [29]. Mean annual precipitation ranges from less than 500 mm in the South of Portugal up to 3000 mm in the North, with 40% of the annual amount falling in winter [26]. High temperatures and high evapotranspiration lead to higher average water consumption per hectare in southern European countries [30]. Water abstraction for irrigated agriculture has changed the flow regime of many river basins and lowered groundwater levels, particularly in these European regions [12]. Episodes of water scarcity occur in PTRH5 (Tejo and Ribeiras do Oeste) and in all RBDs to the south of it where rain is more concentrated in fewer days during the year, and the “normal” year corresponds closely to a “dry” year [31]. Nevertheless, if there is suitable sustainable water management, the available water resources are sufficient to satisfy the needs [32].

The most represented land use classes in Portugal are agricultural areas and forest and semi-natural areas, namely transitional woodland-shrub [33]. The proportion of extensive crop production and extensive grazing is particularly high (59%) which could lead to better ecosystem condition under reduced pressures [34]. Therefore, changes in the vegetation cover and a poorly planned and managed forests have considerable impacts on the water cycle causing pressures on the water environment [12]. Portugal is one of the European countries with the highest diversity of organisms and farming systems, but at the same time is one of the countries more vulnerable to the loss of that diversity [32].

Portugal is also one of the European countries that are reliant on water resources that originate outside their territory [30]. The share of Portugal in the respective international RBDs with Spain is PTRH1 (Minho 9% and Lima 47.6% of the ES010), PTRH3 (19.1% of the ES020), PTRH5A (31.1% of the ES030) and PTRH7 (17.2% of the ES040) [31]. *Fig 1* shows the transboundary water management by RBDs. The boundaries of the National and International RBDs are displayed using version 1.4 of the Water Information System for Europe River Basin Districts dataset available from the European Environment Agency [35]. This dataset is based on data reported to WISE by the EU Member States [35].

**Fig 1.**
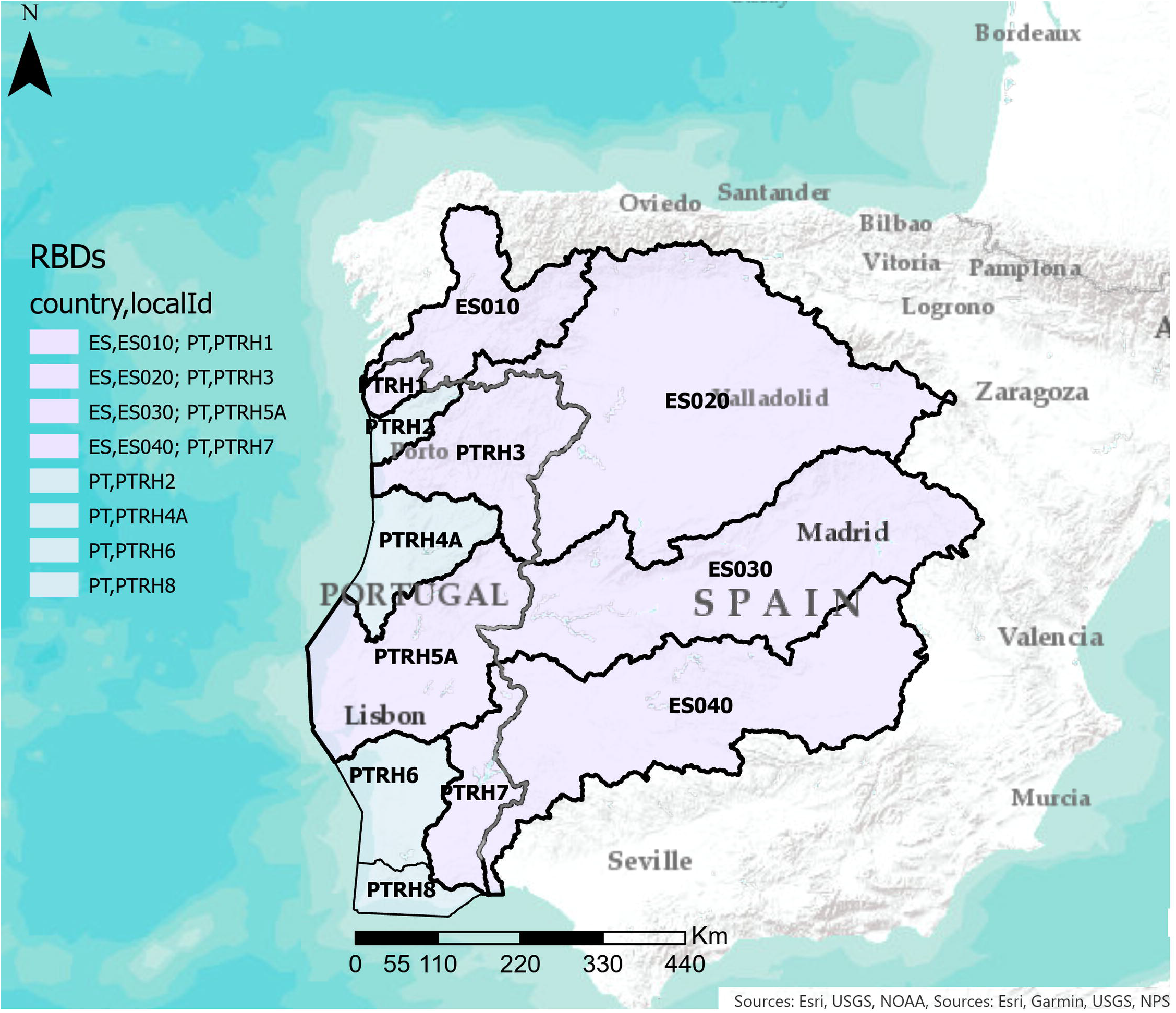
Overview of the national/ international RBDs. The localId refers to the RBDs abbreviations: PTRH1 (Minho and Lima, 2465 km2); PTRH2 (Cávado, Ave and Leça, 3584 km2); PTRH3 (Douro, 19219 km2); PTRH4A (Vouga, Mondego and Lis, 16981 km2); PTRH5A (Tejo and Ribeiras do Oeste, 25665 km2); PTRH6 (Sado and Mira, 12149 km2); PTRH7 (Guadiana, 11611 km2); PTRH8 (Ribeiras do Algarve, 5511 km2); PTRH9 (Azores, 10047 km2); PTRH10 (Madeira, 2248 km2). Source: [35].

At a national level, the main problems associated with the management of water resources are related to water quality, nature conservation, supply and demand, water domain and planning, irrigation, information and knowledge, economic and financial regime, and energy security [32]. Nowadays, hydropower forms a considerable contribution to the Portuguese energy sector representing 30% of the national electricity consumed [35].

According to the most recent European assessment under the Second River Basin Management Plans for Portugal [31], droughts and water abstraction have been reported to be relevant for a major part of Portugal, representing significant pressures on both surface and groundwater ecosystems with direct impacts on water availability. The seriousness of ecosystem degradation and scarcity of the services, as well as the existing institutional context, should always be the starting point for designing analysis and governance solutions [2]. Water yield is a key ecosystem service in river basins [36] and has a provisioning function that represents water balance across space and time [37].

### 2.2. The InVEST Water Yield model

The WYM is defined by physical expressions and spatial and temporal resolutions, providing as outputs maps, outcomes of water services, impacts of environmental changes and analyses of spatial patterns (Sharp et al., 2020). The InVEST WYM approach relates the LULC types with the Evaporative Index (EI) using the methodology developed by [38]. It does the water balance, estimating the annual water yield (Y) for each pixel (x) through the difference between precipitation and actual evapotranspiration. The WYM physical expressions are extensively described in [22] and [38]. The InVEST WYM calculates water yield, water consumption and hydropower valuation. In this study the focus is to calculate freshwater resources availability, i.e. water yield, at a national scale by RBD.

### 2.3. Data

In the InVEST WYM, the runoff from each pixel cell in a catchment is determined based on the water balance concept [39]. For that, the model requires five biophysical parameters as georeferenced rasters (LULC, root depth, plant available water content (PAWC), annual precipitation and evapotranspiration), a vectorial format layer delimiting the study catchment areas, and a biophysical table in CSV format [38].

All required data were obtained freely on the Internet from research centres and public institutions. The datasets Root restriction layer depth and PAWC were provided by the European Soil Data Centre (ESDAC), a European reference Centre for soil-related data [40]. The annual precipitation was obtained from WorldClim database [41] and the reference evapotranspiration from the Consortium of Spatial Information, Global-Aridity and Global-PET Database [42], a product derived from the WorldClim global data. These datasets were modelled based on a high number of climate observations and Shuttle Radar Topography Mission (SRTM) data. The other source of climatic data was obtained by the National Information System of Hydric Resources [43] who collects, processes, and publishes meteorological data at the national scale.

LULC data for years (1990, 2000, 2006, 2012 and 2018) were obtained from Copernicus Global Land Service products [44]. Land cover maps represent spatial information on different types (a total of 44 classes) of physical coverage of the Earth’s surface, using a Minimum Mapping Unit (MMU) of 25 hectares (ha) for areal phenomena and a minimum width of 100 m for linear phenomena.

The CSV biophysical table required by the model has values associated with each LULC class, rooting depth (mm), and plant evapotranspiration coefficient (*k*_*xj*_). This latter is used to calculate potential evapotranspiration by using plant physiological characteristics to modify the reference evapotranspiration. The table was built using data available in the literature [38,45,46].

### 2.4. Sensitivity analysis

Sensitivity analysis was performed by adapting the methodologies proposed by: [15,22,24].Two different data sources of precipitation (P1; P2) and evapotranspiration (E1; E2) were tested. P1 and E1 refer to precipitation and evapotranspiration indices based on satellite data from WordClim database; P2 and E2 were collected, processed, and published by the National Information System of Hydric Resources (SNIRH). The model was also tested with two different PAWC. The main PAWC is the official dataset provided by the European reference Centre for soil-related data. In mainland Portugal this dataset has pixels values varying between 0.03 and 0.14 with a mean value equal to 0.08. The lowest values are mostly related with coarse-textured soils (sands and loamy sands) which have low ability to retain water [47]. Apart from particle size affecting the volume of water that can be stored, different degrees of PAWC fractions depend on soil depth [48]. To test the model under extreme values of PAWC and soil depth, a new dataset was created by reclassifying the original PAWC, obtaining fractions between 0.03 and 0.98. The reclassification was meant to test the model to the maximum possible value, which is 1.

The sensitivity of the model to Zhang seasonality coefficient can also be interpreted as the sensitivity to the PAWC since these parameters play a similar role in the model structure [24]. The Z coefficient (Z1, Z2) is estimated at 0.2*N, where N is the number of rain days (> 1 mm) per year [38]. The N1 values were obtained from the *Portal do Clima* PROJECT [49] which provides several climatic indicators aiming to quantify the occurrence and risk of different atmospheric events [50]. The N2 values were obtained from the Portuguese Institute of the Sea and the Atmosphere [51] database [27]. This institution supports the nation with wide climate-related information products. Table 1 shows the assigned values to this seasonality term (Z) for mainland Portugal.

**Table 1.** The number of rain days (N1, N2) per year and the values assigned to the Zhang coefficient (Z = 0,2*N).

| Year | N1 [49] | Z1 | N2 [51] | Z2 |
| --- | --- | --- | --- | --- |
| 1990 | 82 | 16 | 99 | 20 |
| 2000 | 106 | 21 | 115 | 23 |
| 2006 | 100 | 20 | 102 | 20 |
| 2012 | 94 | 19 | 129 | 26 |
| 2018 | 93 | 18 | 123 | 25 |

Considering the increasing of fire events in the last years [2] and assuming that they have impacts on water ecosystem functions due to loss of diversity and modifications on vegetation structure [52], an alternative biophysical table was created after changing the value of the plant evapotranspiration coefficient (kc) of the LULC classified as 334 (burnt areas) by 10% higher. Uncertainties on this parameter are large since it remains difficult to provide accurate estimates of the actual evapotranspiration from forests [24]. Table 2 summarizes the input data and their sources by the sensitivity analysis tests.

**Table 2.**
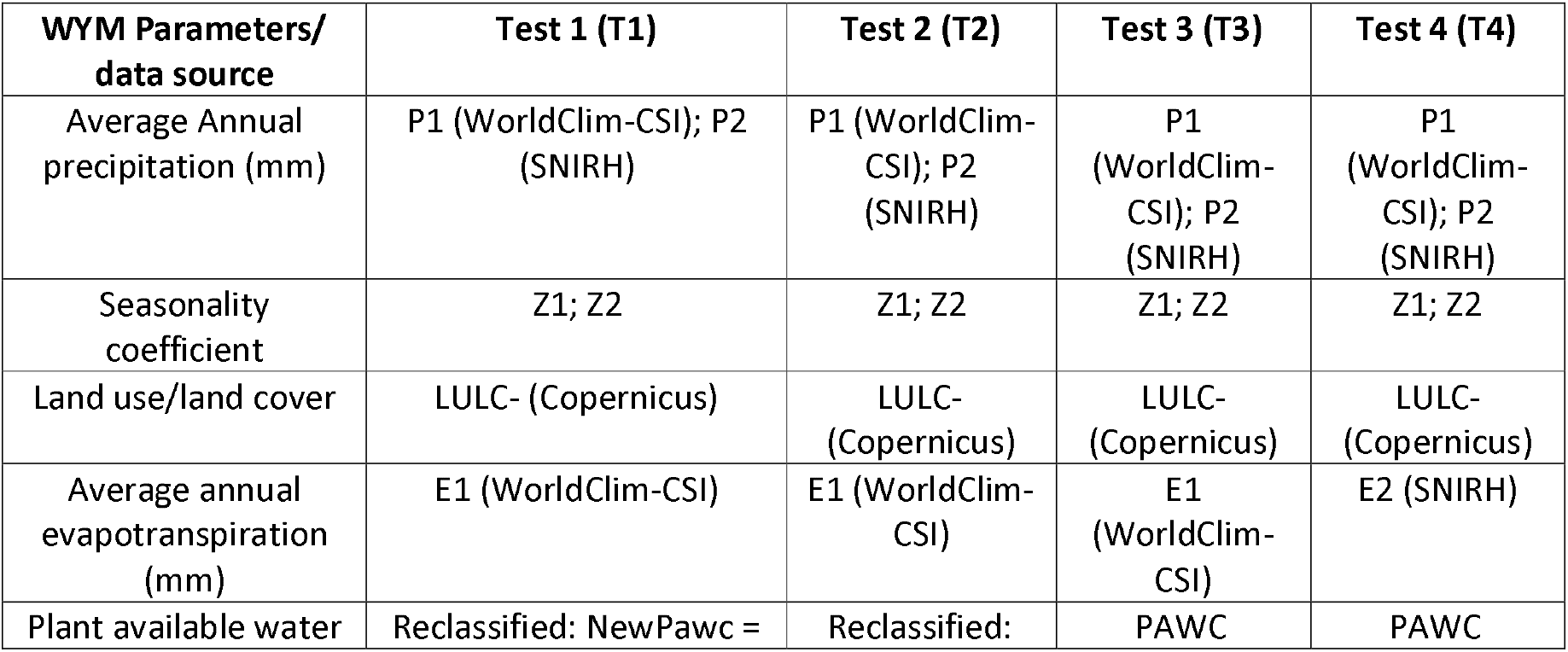
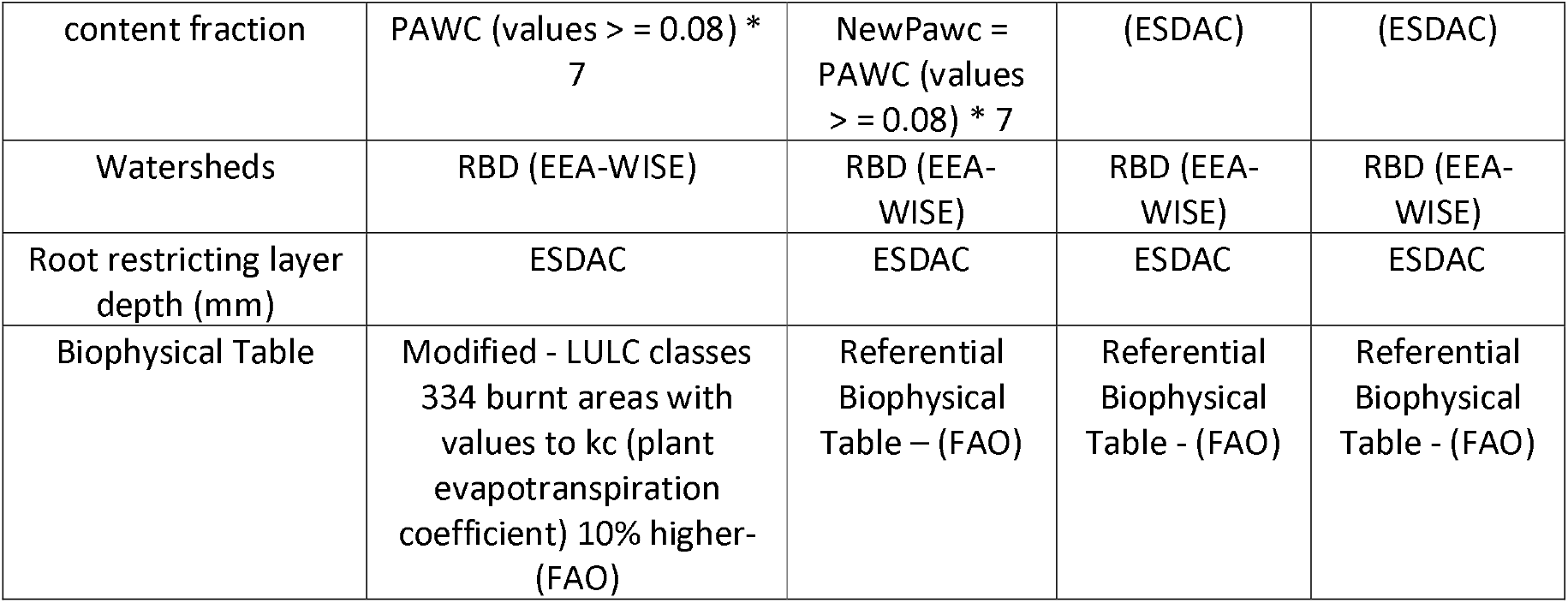
Input data required in the WYM grouped bythe 4 sensitivity analysis tests.

### 2.5. Validation

The minimum requirement when validating assessment models is to establish an adequate scientific basis for credibility [18]. The model is said to be validated if its accuracy and predictive capability in the validation period have been proven to lie within acceptable limits [53]. If a test determines that a model does not have enough accuracy for any of the sets of experimental conditions, then the model is invalid. However, determining that a model has sufficient accuracy for numerous experimental conditions does not guarantee that a model is valid everywhere in its applicable domain [54].

Model validation is defined as the process of demonstrating that a given site-specific model can make sufficiently accurate estimations [53]. The confidence-building view defines the validation of impact assessment models as the process of building scientific confidence in the methods used to perform such an assessment [18].

In this study, is reported the water yield estimates for the 8 RBDs (PTRH1 to PTRH8) covering all mainland Portugal’s extent. These districts represent the administrative jurisdictions responsible to manage the Water Framework Directive (WFD) on the national scale. To validate the model, we compare the mean estimated water yield in the RBDs to the European Environment Agency (EEA) databases on the quantity of Europe’s water resources. The EEA’s Waterbase [35] contains data on freshwater resources availability, delivered by EEA member countries, in the scope of the current WISE SoE - Water Quantity (WISE-3) Water Information System for Europe (WISE).

## 3. Results and discussion

### 3.1. Meteorological data analysis

In the WYM, precipitation and actual evapotranspiration are related to the spatial distribution of meteorological factors and land cover types [39]. To study the meteorological outcomes, we started performing an exploratory data analysis of the precipitation data based on their relationship to some critical output values.

The boxplots in **Fig 2**, show the annual average precipitation by river basin district (RBD). The value is maximum (1850 mm) in the RBD (PTRH1) located in the northwest coast of the country, where the high elevation in the coastal zones contributes to the development of convective systems that promote the occurrence of precipitation events [27]. The minimum value (less than 500 mm) is in the PTRH8 district in the south of Portugal, where are located the most vulnerable regions to drought events, being observed the lowest precipitation indices and the highest temperatures [31].

**Fig 2.**
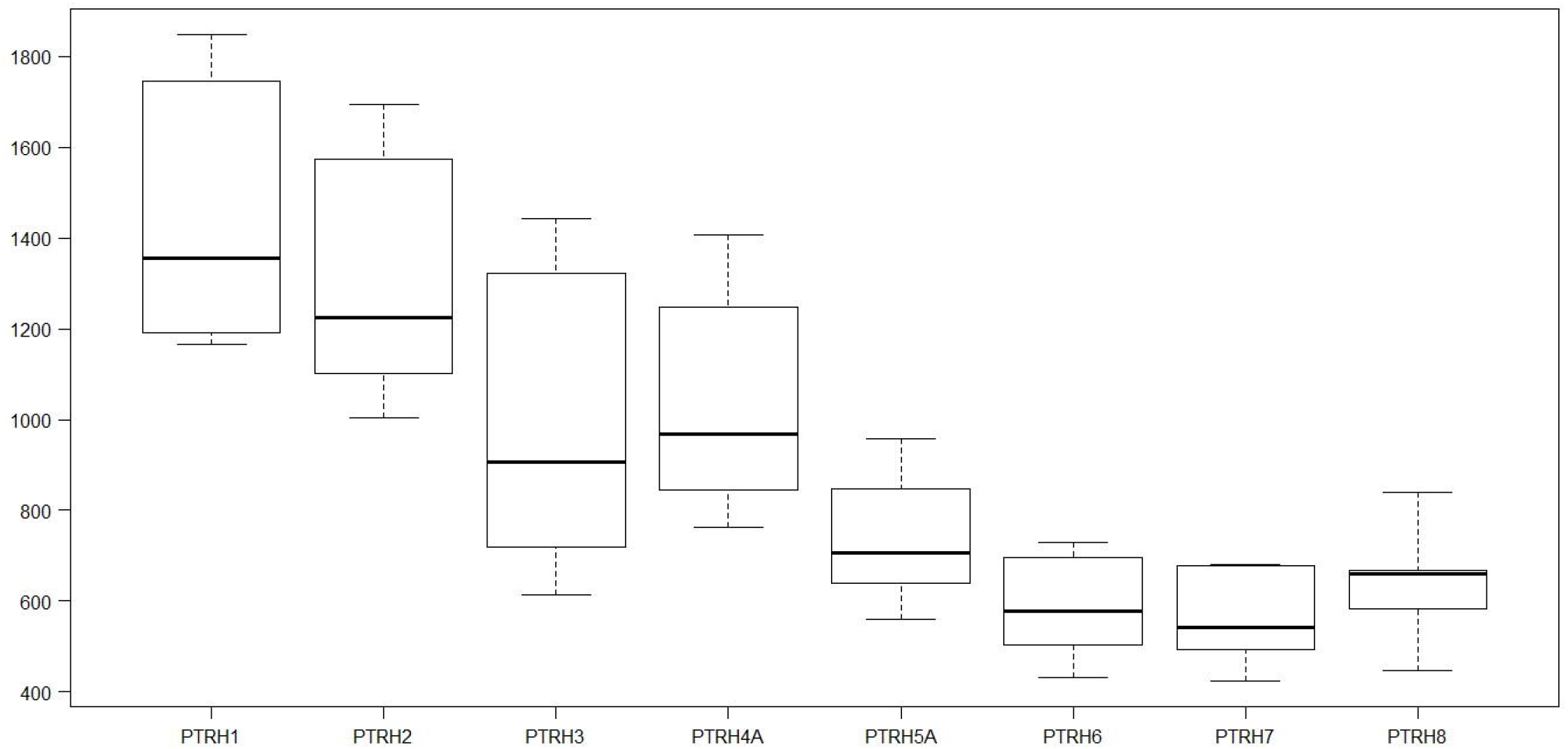
Boxplots: Annual average precipitation by river basin district (mm).

This exploratory analysis is suggesting two main patterns, one with the highest precipitation indices and other with the lowest. Through a spatially exploratory analysis in a GIS environment, the study area could be divided into two sectors, following the concept that Portugal is divided between North and South by the Tagus river that cuts the territory East to West [55].

The north of the country (NPT) is representing 40.62% of the total study area, including the RBDs: PTRH1, PTRH2, PTRH3, PTRH4; and the other sector (SPT – South of Portugal) is representing 59.38% and includes the following river basin districts: PTRH5, PTRH6, PTRH7, PTRH8.

**Fig 3** is showing plots of the annual average precipitation (P1 – WorldClim; P2 – SNIRH) in the study years (1990, 2000, 2006, 2012 and 2018), by these 2 sectors, P1, P2 for NPT (North of Portugal) and P1, P2 for SPT (South of Portugal).

**Fig 3.**
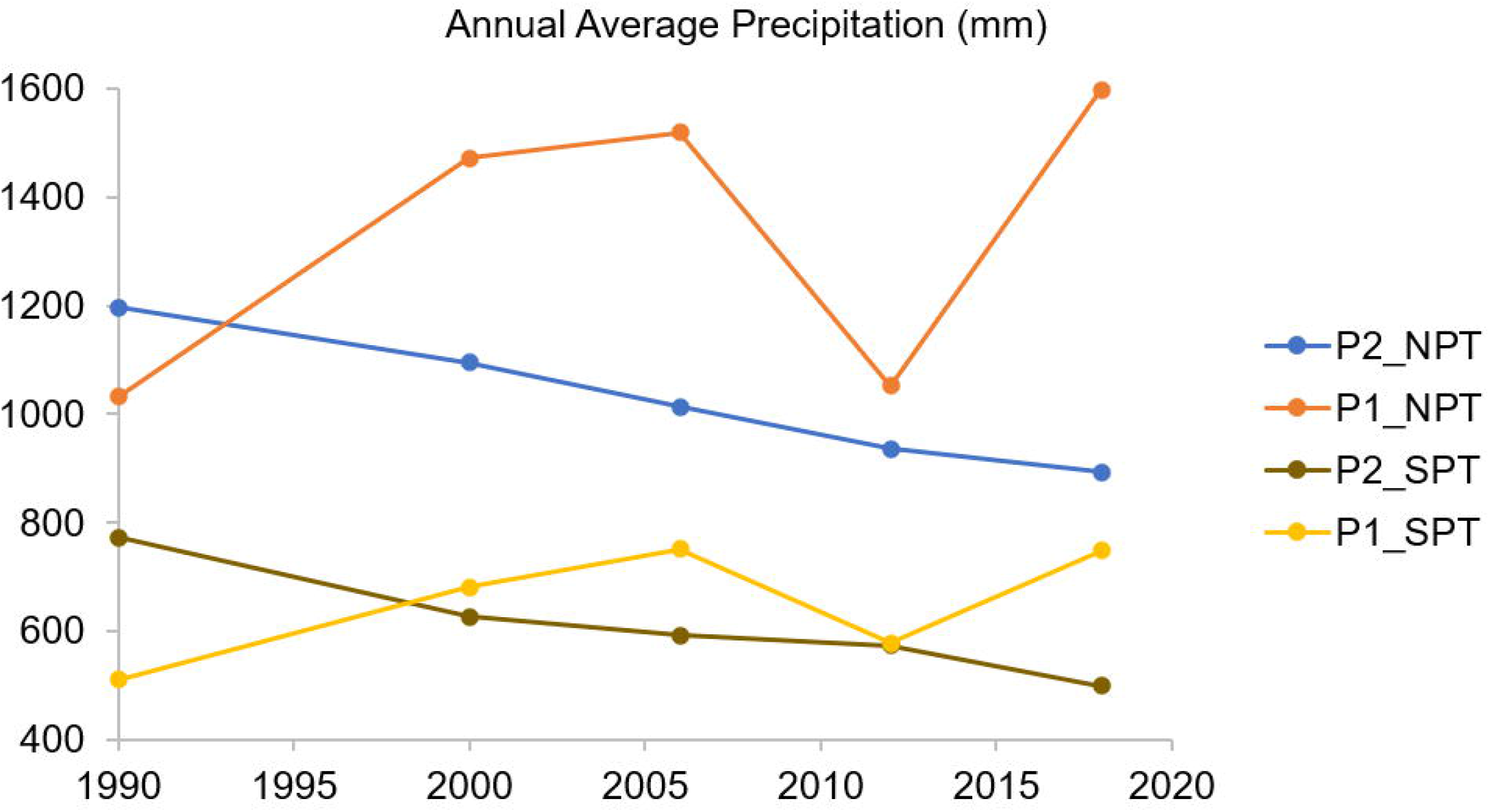
Annual average precipitation (mm) by data source and sectors. P1 – WorldClim; P2 – SNIRH; NPT – North of Portugal (PTRH1, PTRH2, PTRH3, PTRH4); SPT – South of Portugal (PTRH5, PTRH6, PTRH7, PTRH8).

The precipitation data from SNIRH (P2) shows a continuous decreasing of the values over the years, while the data from WordClim (P1) shows high variability. Pearson’s correlation between P1 and P2 in the NPT sector is equal to -0.409, in the SPT sector is -0.754. The spatial variability from North to South of Portugal is explained by the latitude and topographic configuration that influence the rain events and their temporal distribution [27]. As referred by [26], the years 1990 and 2012 are identified as being particularly dry within the Iberian Peninsula territory recording severe drought events.

### 3.2. Dryness Index & Evaporative Index outcomes

The Budyko curve relates to the Dryness Index (DI) and the evaporative index (EI). Together, these indexes define the energy limit and the water limit of a hydrologic system. The DI is also applied to derive climate classifications for different zones: Hyper-arid (DI > 20), Arid (5 < DI ≤ 20), Semiarid (2 < DI ≤ 5), Dry sub-humid (1.5 < DI ≤ 2) and Humid (DI < 1.5) [56]. The EI values close to 1 mean that almost all amount of rain is returning to the atmosphere by evapotranspiration from the vegetation and soil, and the lowest values indicate higher water yield capacity.

The University of Leuven, with support of the International Water Management Institute and the International Centre for Integrated Mountain Development [42] estimated for mainland Portugal an area of 46365 km^2^ that is highly vulnerable to drought events and is classified as an arid zone. This area represents 50.36% of the country’s extent, which is comparable with the sector (SPT – South of Portugal) that represents 59.38%.

The mean difference between the global reference evapotranspiration (E1) and that obtained from the national database (E2) is nearly -10%. E1 shows a standard deviation equal to 31 mm against 2.5 mm in E2, these values were obtained comparing datasets from years (1990, 2000, 2006, 2012 and 2018). The following boxplots are representing the DI **(Fig 4)** & EI **(Fig 5)** results by RBD. It is showing the same pattern occurred with the precipitation data permitting to group the national river basin districts in two sectors, NPT – North of Portugal (PTRH1, PTRH2, PTRH3, PTRH4) and SPT – South of Portugal (PTRH5, PTRH6, PTRH7, PTRH8).

**Fig 4.**
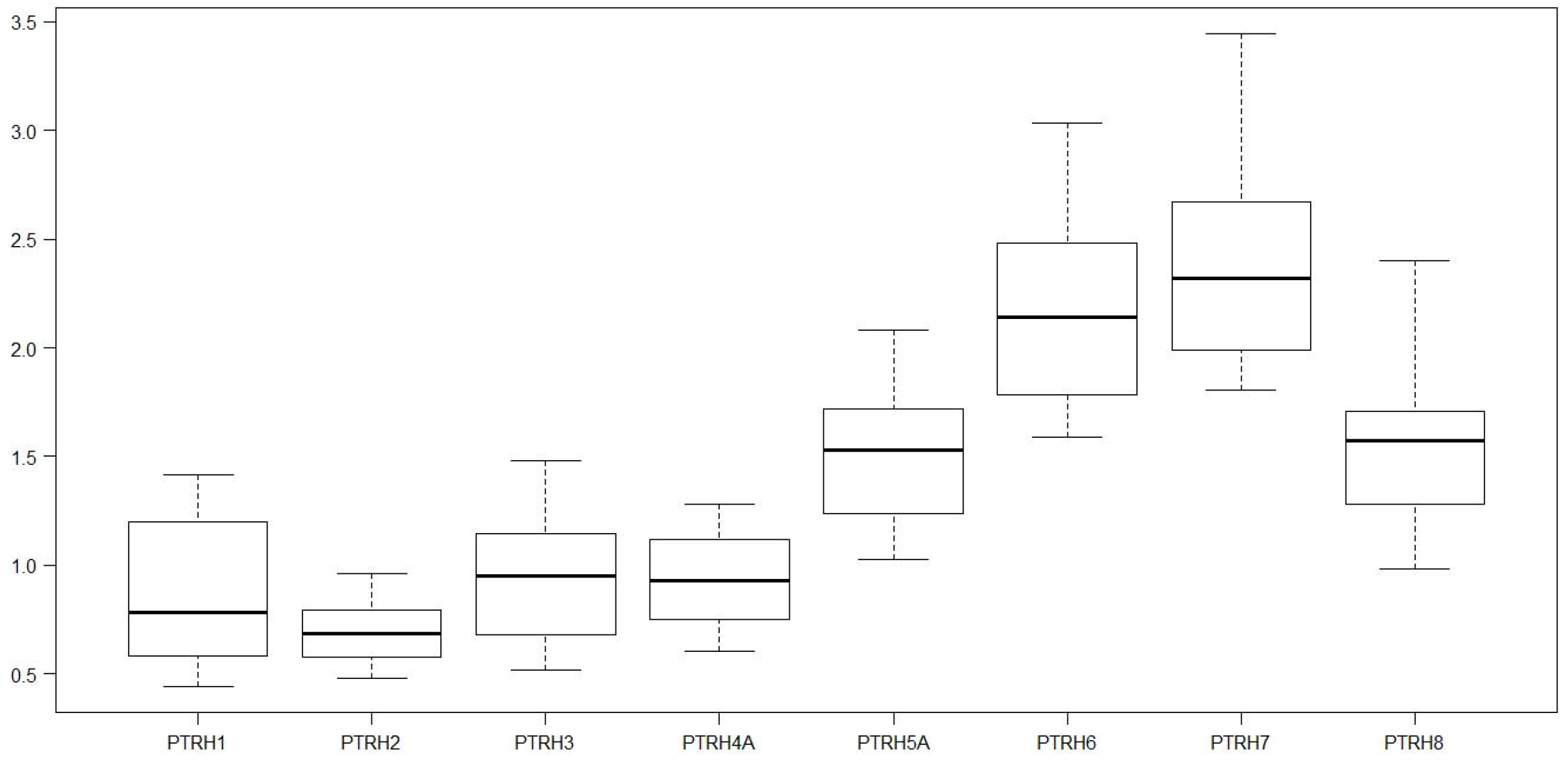
Boxplots: Dryness indexes by RBD.

**Fig 5.**
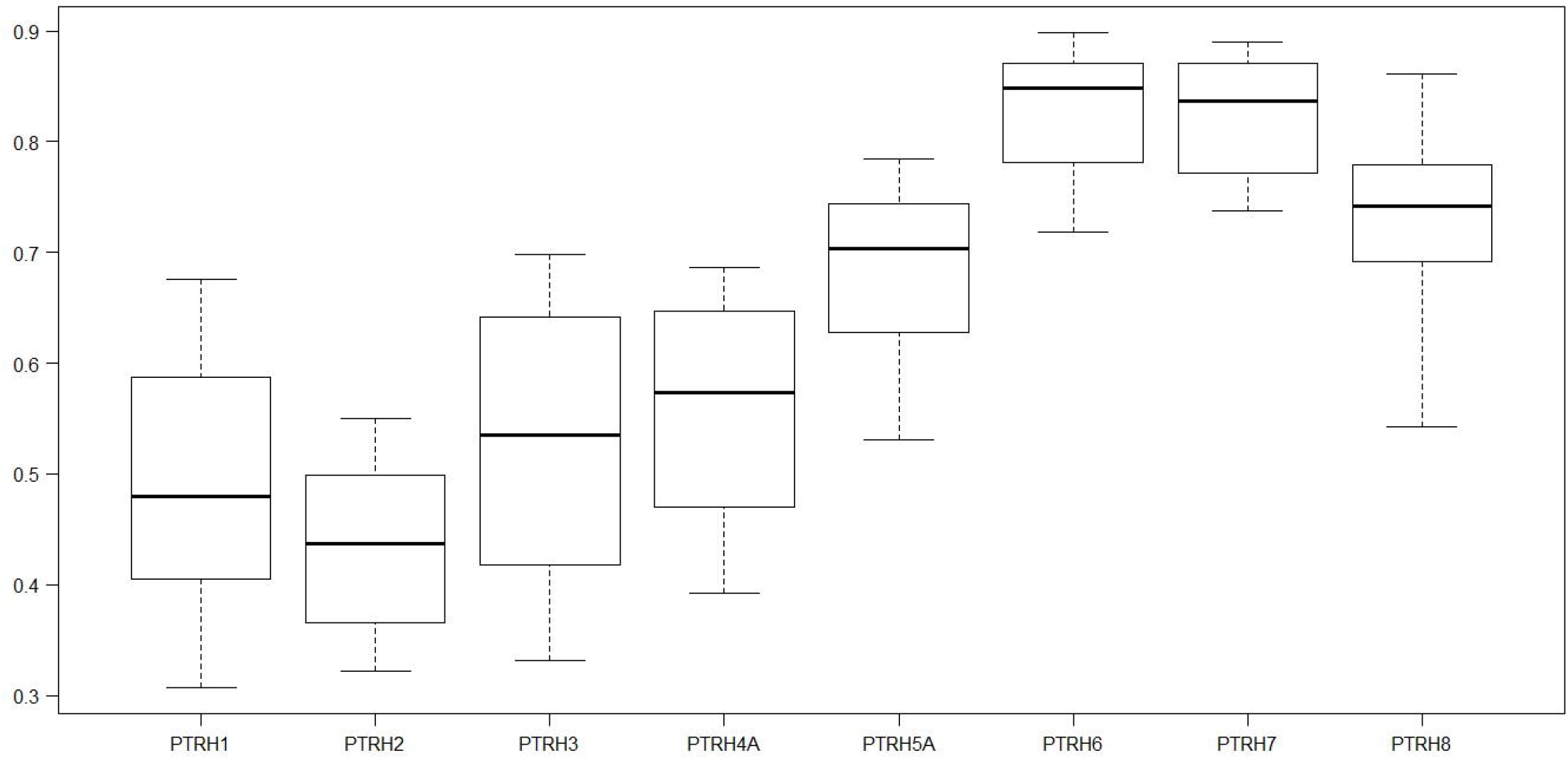
Boxplots: Evaporative indexes by RBD.

The highest DI & EI values are in the SPT sector. These values are showing the high vulnerability of this sector to severe drought events. The EI values classify SPT sector in Dry sub-humid and Semiarid zone and the NPT sector as Humid/Dry sub-humid zone as proposed by the Global Map Aridity [42].

The Budyko curve is bounded by two limits – an energy limit in which actual evapotranspiration (AET) is equal to the potential evapotranspiration, and a water limit for which actual evapotranspiration is equal to precipitation [38]. Due to spatial and temporal variability in climate forcing, the asynchronicity of water availability (P) and demand (PET), the imperfect capacity of the root zone to buffer that asynchronicity, and lateral redistribution of water within the catchment, the Budyko curve lies below those two limits [24]. To understand the concept and to verify the WYM, a Budyko curve graphic with DI & EI outcomes is plotted in Fig 6. It shows results of the sensitivity analysis simulations for tests T3 (E1) and T4 (E2).

**Fig 6.**
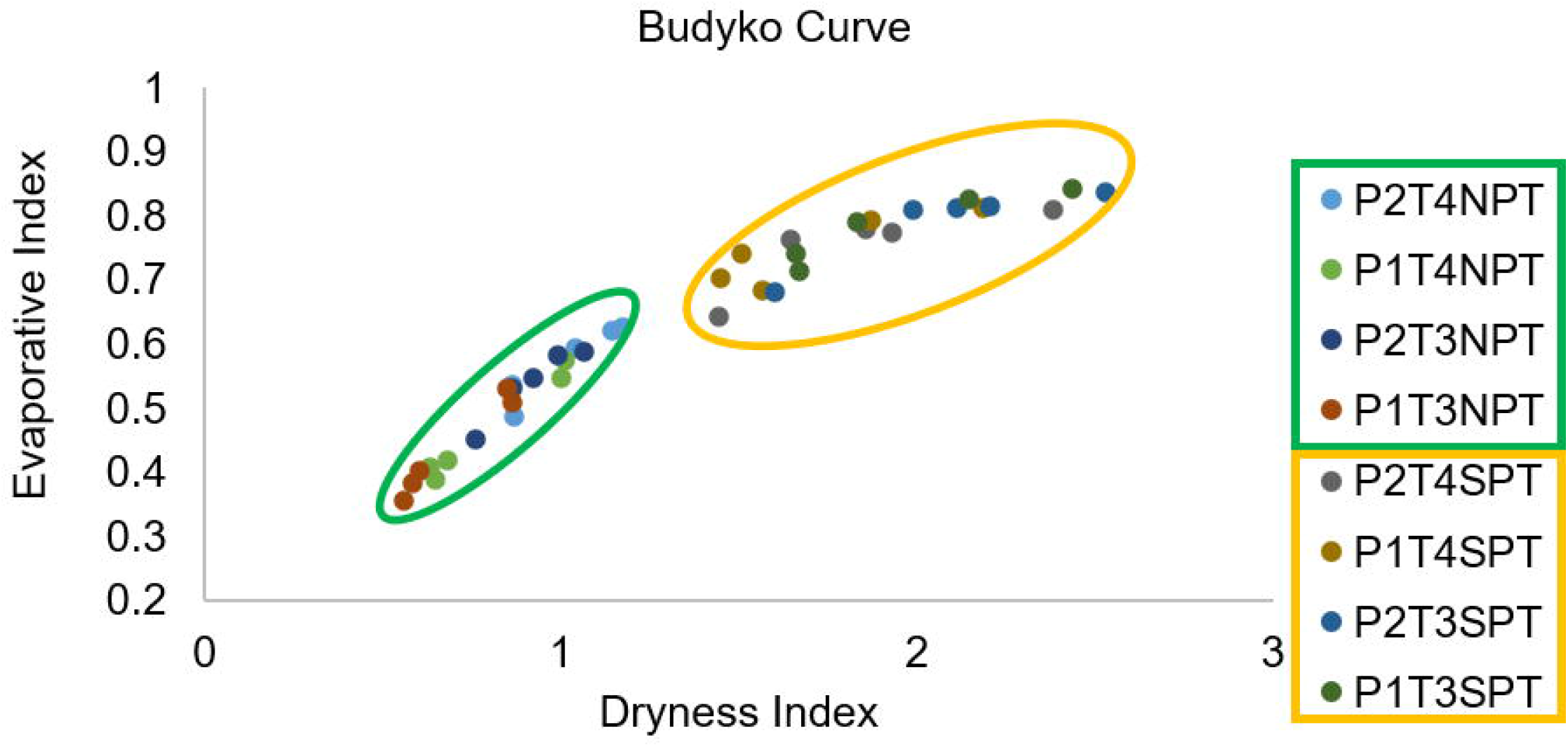
Budyko curve by sensitivity analysis tests. P1 – WorClim; P2 – SNIRH; T3 – E1; T4 – E2; Green Cluster: NPT (PTRH1, PTRH2, PTRH3, PTRH4); Yellow Cluster: SPT (PTRH5, PTRH6, PTRH7, PTRH8).

Examining the Budyko curve with mean values of DI and EI of the five studied years, we can distinguish two main clusters referring to these two sectors, NPT (North of Portugal) and SPT (South of Portugal). As water yield is greater in those points close to zero, the cluster formed by points from the NPT sector show better water availability, with the greatest value in the simulation T3 using P1 and E1, and for the SPT sector, with test T4 using P2 and E2.

### 3.3. Corine LULC changes

The change in land use can influence the integrity of natural systems [8] and the WYM is assessing the impacts of land cover changes in each pixel cell offering insights into how changes in land-use patterns affect annual surface water yield [38]. The five main categories of the Corine land cover datasets are artificial surfaces (1), agricultural areas (2), forest and semi-natural areas (3), wetlands (4) and water bodies (5).

Analyzing the land cover changes in national scale (LULC_PT) in the five main Corine Land Cover categories between 1990 and 2018, the numbers show that artificial areas have increased more than 100%, followed by surface water bodies and wetlands, with 34% and 7%, respectively. Agricultural, and forest and semi-natural areas have decreased by 2% and 3%, respectively. The studied categories followed the same pattern in the NPT and SPT sectors (LULC_NPT; LULC_SPT).

Fig 7 shows the evolution (CLC_Evolution) and variation (Var_18_90) for the main land use/land cover categories of Corine data (1-Artificial surfaces, 2-Agricultural area, 3-Forest and seminatural areas, 4-wetlands, 5-water bodies). Each red dot is referring to values estimated for the studied years (1990, 2000, 2006, 2012, and 2018). The variable Var_18_90 is the variation between years 1990 and 2018.

**Fig 7.**
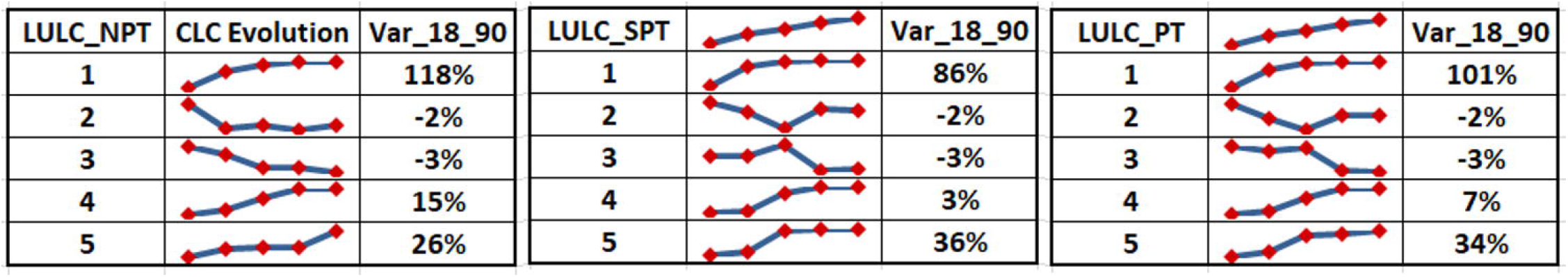
Tables showing Corine Land Cover evolution (CLC_Evolution) and variation (Var_18_90) for the main land use/land cover categories. 1-Artificial surfaces, 2-Agricultural area, 3-Forest and seminatural areas, 4-wetlands, 5-water bodies. Each red dot is referring to values estimated for the studied years (1990, 2000, 2006, 2012, and 2018). LULC (Land Use/Land Cover), NPT (North of Portugal), SPT (South of Portugal), PT (mainland Portugal).

At the national scale, as the number of artificial areas increased the number of superficial water bodies also increased, showing a Person’s correlation equal to 0.897. The inverse situation is observed when comparing artificial areas with agricultural and forest areas, showing a Person’s correlation equal to -0.702 and -0.661, respectively.

The growing of the artificial areas demands an effort of the water-related ecosystems services. To respond to that, in the last 18 years, many dams have been constructed, to satisfy water supply, agriculture irrigation, industry and to deal with water scarcity in the most vulnerable regions. The Alqueva dam, located in the SPT sector is the largest superficial water body in Europe [35], and it started being operational in 2002 in the Guadiana River.

### 3.4. Sensitivity analysis

Sensitivity analysis provides a logical and verifiable method of optimizing the distribution of resources used to determine the most important parameters [18]. Considering the spatial pattern established by the meteorological outcomes, this step began investigating the sensitivity of the WYM outcomes by sectors NPT (North of Portugal) and SPT (South of Portugal). We assessed and compared the model’s performance over sensible ranges to determine the effect of these variations. Fig 8 shows water yield result of each test by RBD for NPT sector. The year 2006 has only 8 estimated WY values because Z1 was equal to Z2.

**Fig 8.**
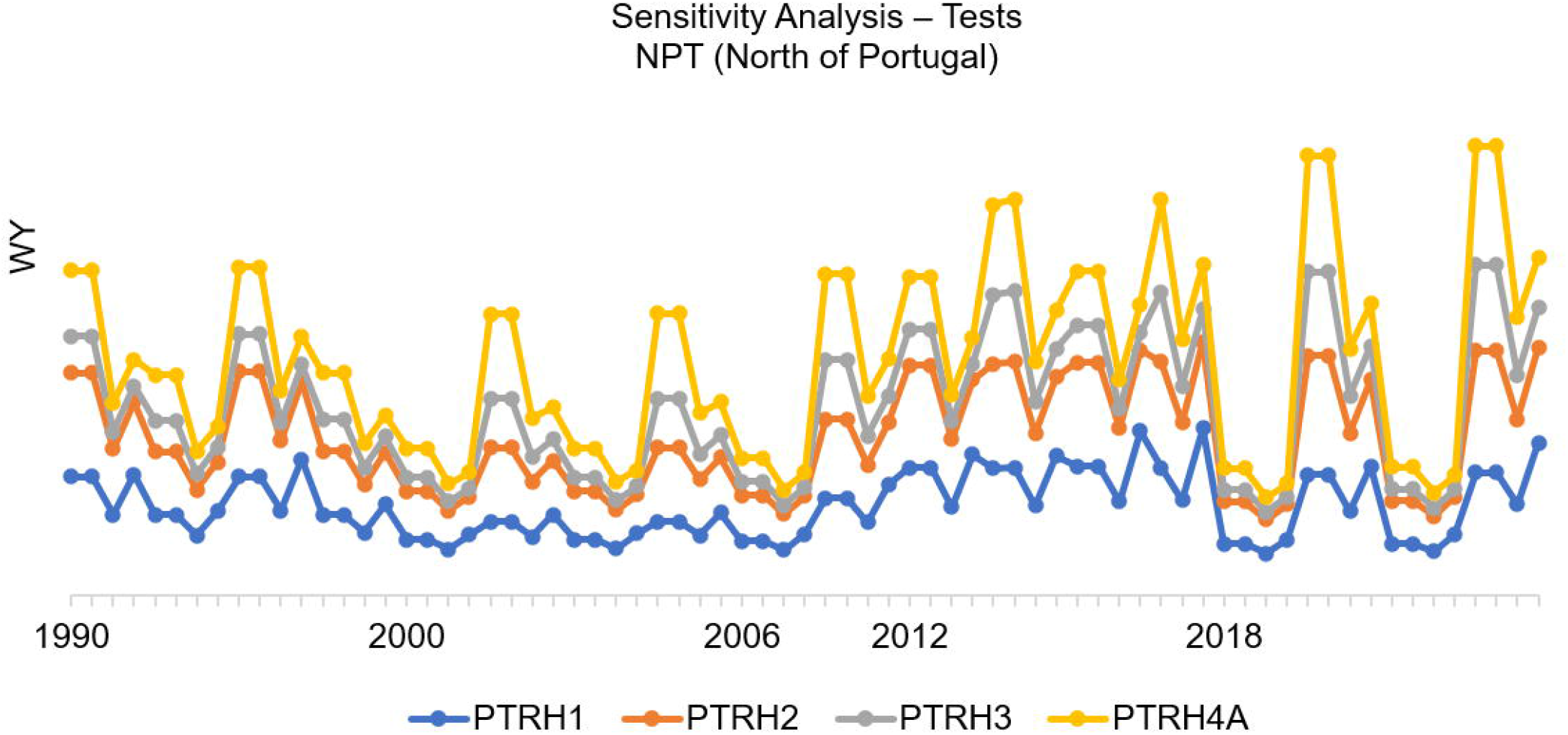
Sensitivity analysis tests by river basin district (RBD) of the North of Portugal.

In the NPT sector, 91% of the simulations show a Person’s correlation of more than 0.85, against 0.58 in the SPT sector (Fig 9). The lowest correlation observed on both sectors was 0.6. The NPT sector shows sensitivity to all the tests throughout the studied years. The SPT sector shows high sensitivity, especially in the years 1990, 2012 and 2018. In most of the tests, when using the largest value to Z (Z2), the results are slightly higher when compared to Z1. The PTRH5A shows the lowest variations between tests.

**Fig 9.**
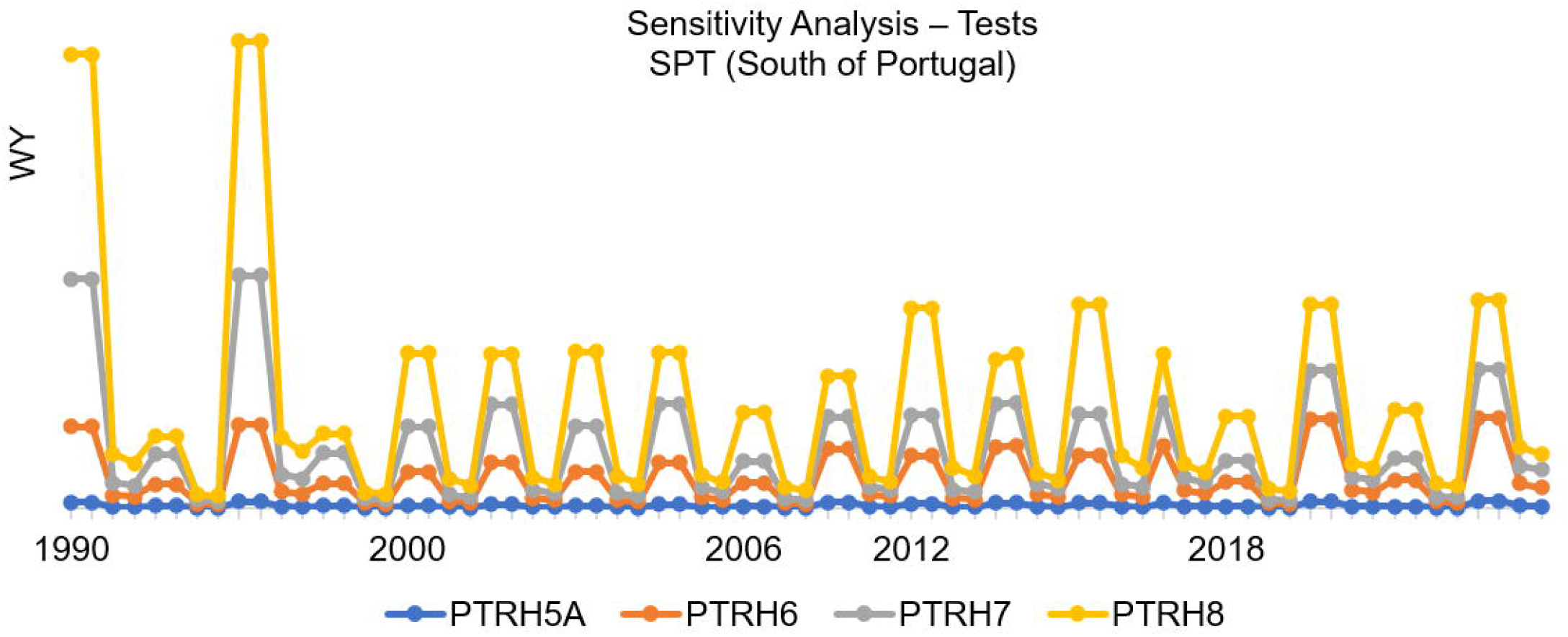
Sensitivity analysis tests by river basin district (RBD) of the South of Portugal.

The RBDs in the SPT sector show higher sensitivity to the alternative PAWC (T1 and T2). In both sectors, the model shows higher sensitivity to the tests in the years 1990 and 2012, where is known that the annual precipitation was lower due to the drought events occurred in those years, as referred in the literature [26]. The storage capacities provided by reservoirs and lakes drive the possibility for water to be spared between periods with recharge (high precipitation indices) and periods of consumption [57]. Catchments with high resistance can store water over long periods (months or years) and release water gradually to the stream [13] that can also affect the results.

The highest sensitivity in the PTRH7 is observed when running the model with an alternative PAWC (T1, T2) and alternative biophysical table (T1). Those extremes values have occurred because in this RBD exists the largest superficial water body in Europe, the Alqueva dam [31], and the PAWC in those pixels had a maximum possible value which is 0.98. The reservoir behaves as a “tank” receiving water from rainfall on the surface and inlets or returns and exporting water through evaporation, infiltration, losses, withdrawals and outlets [57].

### 3.5. Model validation

The most convincing evidence that a scientific theory or model is correct is through direct comparison of model predictions with experimental observations [18]. The EEA’s Waterbase [35] contains the volume of freshwater resources by RBD for 2018, that will be compared with the best simulations for the same year.Fig 10 presents the estimated volume of water yield in 2018 by RBD of the North of Portugal and the observed volumes (WISE18). The best performance simulation in this year was Z2P2T4 which is marked in the legend of the image.

**Fig 10.**
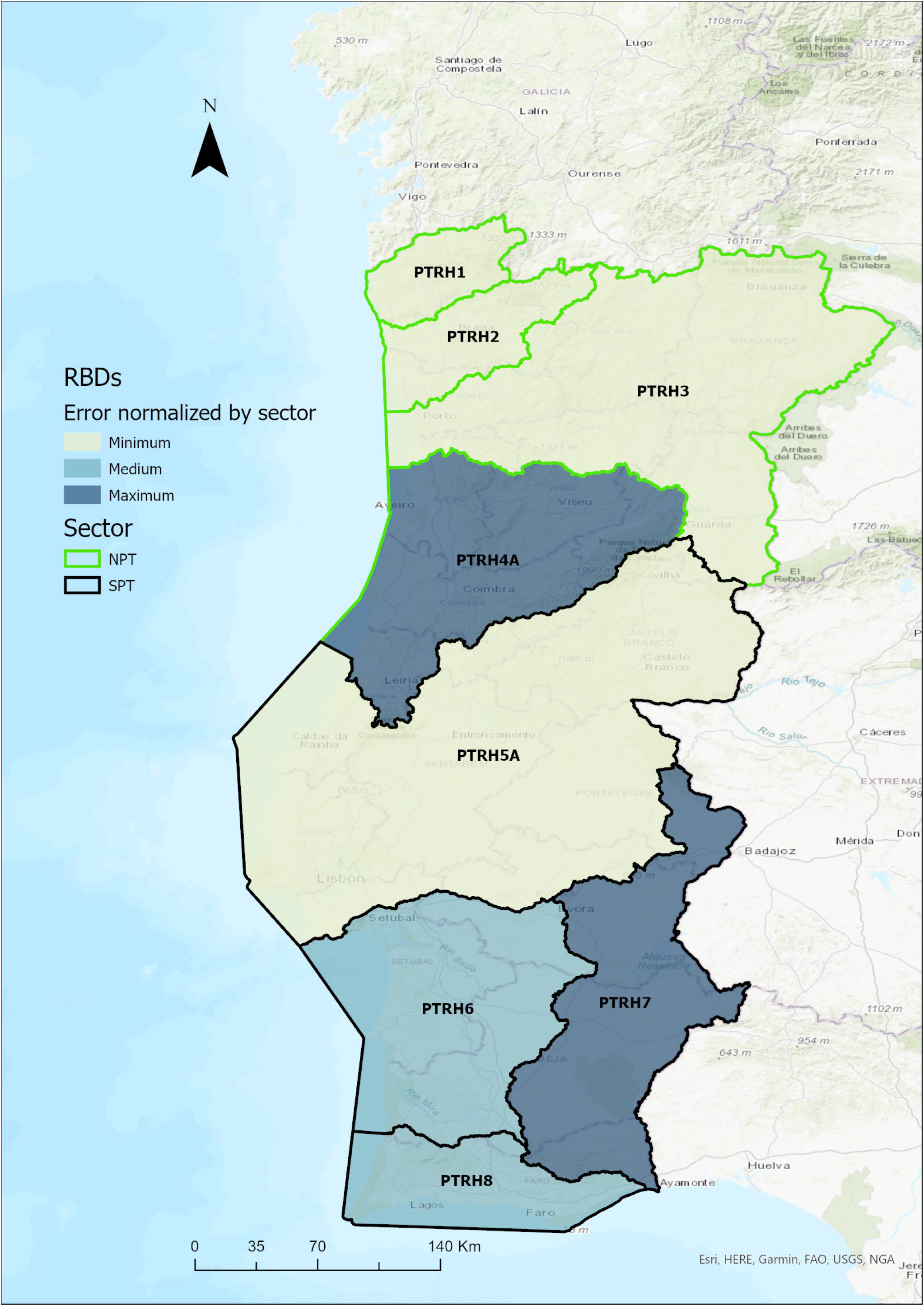
Estimated WY volume (m3) in 2018 by RBD/NPT sector and observed values.

In the SPT sector (Fig 11) the best simulation was considering the test Z1P1T4. The differences between these two tests are on the number of rain days considered, and the source of the precipitation data.

**Fig 11.**
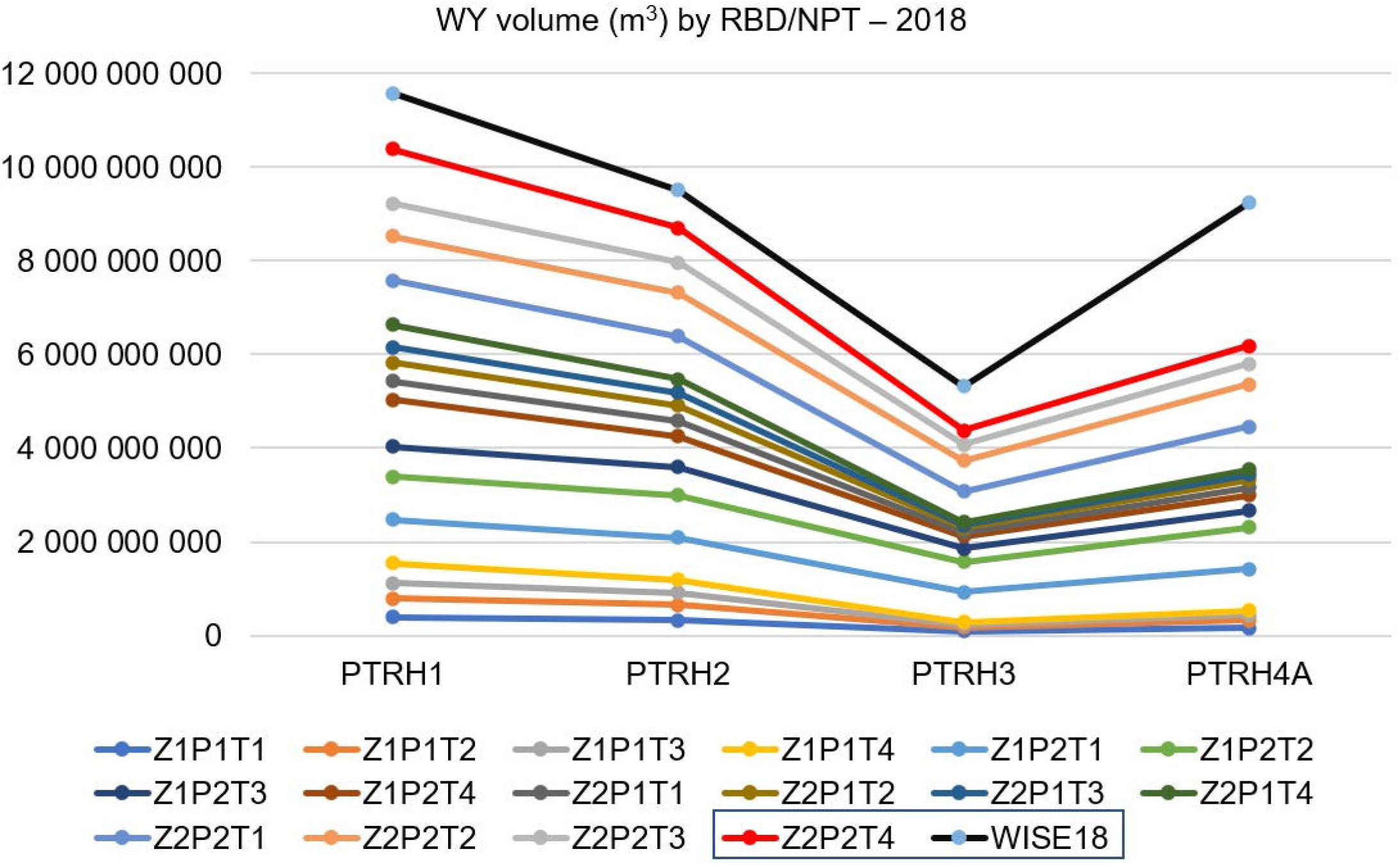
Estimated WY volume (m3) in 2018 by RBD/SPT sector and observed values.

After calibrating the model by sectors with the best simulation performance in 2018, the mean volume of the estimated water yield by RBD was compared with the freshwater resources availability volume, published by [35]. Results at the national level show a correlation coefficient of 0.803 with statistical significance for 0.01 one-tail. Table 3 has the values used to estimate model’s accuracy.

**Table 3.** Mean estimated WY, observed values and the error in the estimations.

| Sector | NPT |  |  |  |  | SPT |  |  |
| --- | --- | --- | --- | --- | --- | --- | --- | --- |
| Best performance/sector | Z2P2T4 |  |  |  |  | Z1P1T4 |  |  |
| RBD | PTRH1 | PTRH2 | PTRH3 | PTRH4A | PTRH5A | PTRH6 | PTRH7 | PTRH8 |
| area (ha) | 246500 | 358400 | 1921900 | 1698100 | 2566500 | 1214900 | 1161100 | 551100 |
| WY_Est_2018 (m<sup>3</sup>/ha) | 4746 | 2037 | 159 | 224 | 31 | 468 | 449 | 2323 |
| WY_Obs_2018 (m<sup>3</sup>/ha) | 4851 | 2280 | 492 | 1801 | -176 | 1455 | -1368 | 1033 |
| error/RBD (m<sup>3</sup>/ha) | -105 | -243 | -333 | -1577 | 207 | -988 | 1817 | 1290 |
| Mean error/sector | -56.5 mm/year |  |  |  |  | 58.1 mm/year |  |  |

WY\_Est\_2018 (mean estimated WY); WY\_Obs\_2018 (mean observed WY - WISE WaterBase).

The WY in the NPT sector was underestimated by 56.5 mm/year and the SPT sector was overestimated by 58.1 mm/year. The difference in the estimations by sectors is explained by the spatial and temporal variability of precipitation and the sensitivities of the model to the climatic variables. Fig 12 shows the spatially assessment of the errors normalized by each sector. In the NPT sector, aside of the RBD PTRH4A that has the maximum error, all others RBDs show minimum normalized errors. In the SPT sector the maximum error was in the RBD PTRH7, medium errors in the PTRH6 and PTRH8, and minimum normalized error in the PTRH5A.

**Fig 12.**
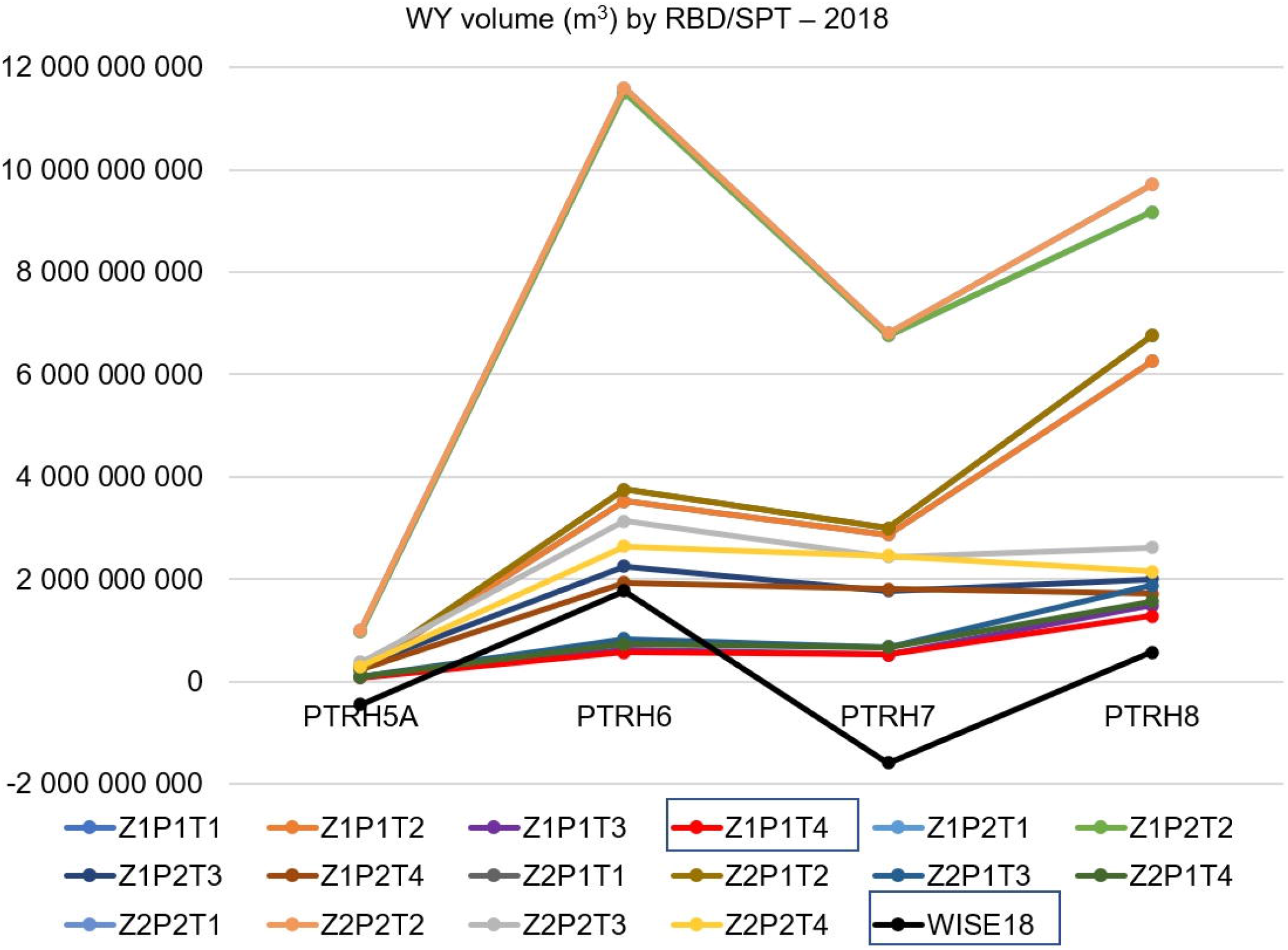
RBD error normalized by the mean error of each sector.

The errors in the estimations can be explained by other variables that the is not taking in account, such as surface runoff, losses or gains to groundwater systems, and storage dynamics. Complex land use patterns or underlying geology may induce complex water balances as it is not precisely captured by the model [38]. However, the InVEST WY model shows a good capability to model water balance and water circulation through sub-surface porous media, and to analyse the effects of meteorological pressures on water ecosystem conditions at RBD level.

## 4. Conclusions

Through the analytical examination of the sensitivity tests, this study contributes to a better understanding of which input parameters affect most the model’s behaviour. The sensitivity of the WYM to the climatic variables was confirmed when comparing simulations with two different climatic datasets, and its sensitivity to land cover changes was confirmed when running simulations with the alternative PAWC. The drought events which occurred in mainland Portugal in the years 1990 and 2012, helped to understand the effects of the meteorological conditions in the freshwater availability. Comparing the model’s performance for the sensitivity tests, the WYM shows to be reactive to changes in the Zhang values (number of rain days in a year), precipitation, and evapotranspiration indices. The model has shown the highest sensitivities when extreme climate conditions occurred.

Models’ accuracy was verified calculating the correlation coefficient at the national level and comparing the mean estimated water yield with mean observed values. Mean WY was underestimated in the NPT sector, and overestimated in the SPT sector. The estimations would be better if the model were calibrated and run separately for each RBD. Nevertheless, the model shows correlation coefficient of 0.803 with statistical significance for 0.01 one-tail. The uncertainties that exist in the model, such as the fact that the WYM is not considering surface water and groundwater interactions and is not differentiating surface runoff and baseflow, combined with complex land-use patterns, such as wetlands, or underlying geology, such as alluvial deposits, could explain the difference between estimated values and observed values.

The comparison of the simulations’ tests and the validation of the model were important to build confidence in the models’ outputs, to support studies related to the spatial estimation of water availability at sub-watershed, watershed, or RBD, and to provide guidance for future research. Better performance can be achieved considering multi-site calibration/validation and multi-parameters checks, especially if spatially variability is observed, and complex land cover and geology are known.

